# A new model explaining the origin of different topologies in interaction networks

**DOI:** 10.1101/362871

**Authors:** Rafael B. P. Pinheiro, Gabriel M. F. Félix, Carsten F. Dormann, Marco A. R. Mello

## Abstract

The architecture of interaction networks has been extensively studied in the past decades, and different topologies have been observed in natural systems. Despite several phenomenological explanations proposed, we still understand little of the mechanisms that generate those topologies. Here we present a mechanistic model based on the integrative hypothesis of specialization, which aims at explaining the emergence of topology and specialization in consumer-resource networks. By following three first-principles and adjusting five parameters, our model was able to generate synthetic weighted networks that show the main patterns of topology and specialization observed in nature. Our results prove that topology emergence is possible without network-level selection. In our simulations, the intensity of trade-offs in the performance of each consumer species on different resource species is the main factor driving network topology. We predict that interaction networks with low species diversity and low dissimilarity between resources should have a nested topology, although more diverse networks with large dissimilarity should have a compound topology. Additionally, our results highlight scale as a key factor. Our model generates predictions consistent with ecological and evolutionary theories and real-world observations. Therefore, it supports the IHS as a useful conceptual framework to study the architecture of interaction networks.

## Introduction

In the past decades, network science, by focusing on the structure of entire systems instead of species, proved to be an outstanding tool for the study of ecological interactions (Dormann *et al.* 2017). One persisting controversy in the literature is the predominant architecture among interaction networks. Two main topologies have been proposed as almost universal: nested and modular (Fortuna *et al.* 2010).

Several studies have detected significant nestedness in interaction networks (Bascompte *et al.* 2003; Guimarães *et al.* 2007b). In a perfectly nested network, the links (i.e., connection between two species in a network) made by species with fewer interaction partners (i.e., other species to which it is connected) tend to be a subset of the links made by species with more interaction partners (Bascompte & Jordano 2007), so interaction overlap is maximum. Nevertheless, several other studies have found a modular topology in interaction networks. A modular network is composed of subgroups of densely connected species (Guimerà *et al.* 2010; Bellay *et al.* 2011; Watts *et al.* 2016).

Contrary to nestedness, modularity is characterized by each node interacting preferentially with a particular subgroup of nodes, overlap is reduced, and several links are considered forbidden (e.g., impossible to occur due to trait mismatch, Jordano 2016). Usually, modules are composed of phylogenetically close species (Krasnov *et al.* 2012) or species that converge in a set of traits (Mello *et al.* 2011). Despite nestedness and modularity being logically different topologies (Ulrich *et al.* 2017) and usually negatively correlated with one another in empirical ecological networks (Thebault & Fontaine 2010; Pires & Guimaraes 2012; Trøjelsgaard & Olesen 2013), networks combining some degree of both have been observed in nature (Olesen *et al.* 2007; Bellay *et al.* 2011; Flores *et al.* 2013).

Diverse explanations to the emergence of each network topology have been proposed For instance, interactions driven by abundance (Vázquez *et al.* 2007), neutrality (Krishna *et al.* 2008), and morphological constrains (Stang *et al.* 2007) for nestedness. And phylogenetic conservatism (Krasnov *et al.* 2012), functional complementarity (Montoya *et al.* 2015), and trait-matching (Donatti *et al.* 2011) for modularity. Interaction intimacy does also seem to play a role in shaping network topology (Hembry *et al.* 2018).

Additionally, a recurrent hypothesis is that nestedness should be expected in mutualisms while modularity should emerge in antagonisms (Thebault & Fontaine 2010). Nevertheless, several studies found empirical evidence against this hypothesis (Olesen *et al.* 2007; Mello *et al.* 2011; Pires & Guimaraes 2012). Despite a diversity of phenomenological explanations, we still poorly understand the mechanisms that drives the establishment of links and so shape network architecture, an issue already pointed out (Ings *et al.* 2009), but which still has not been properly addressed. Maybe as a symptom of this knowledge gap, community-level selection is commonly invoked to explain interaction network topology, despite the strong criticism against it in the evolutionary literature (see Pires & Guimaraes 2012). In the present study, we use a recent hypothesis to propose a unified mechanism that drives the formation of links and scales up to shape network topology.

The integrative hypothesis of specialization (IHS), (early called the integrative hypothesis of parasite specialization, Pinheiro *et al.* 2016; Felix *et al.* 2017), is aimed at explaining the relationship between performance and specialization in consumer-resource interactions (e.g., parasite-host, prey-predator, plant-pollinator). A classical hypothesis states that, due to trade-offs involved in specialization, generalist consumers should be outperformed by specialist consumers in exploiting each resource (Futuyma & Moreno 1988). It is illustrated by the figure of speech “jack-of-all-trades, master of none”. In this scenario, because of those trade-offs, each consumer species tends to specialize in one or few resource species, and several interactions are forbidden. Indeed, some studies have found compelling evidence corroborating this hypothesis in different systems (Poulin 1998; Muchhala 2007). However, other studies found that generalistic consumers achieve higher performance in exploiting each resource (Krasnov *et al.* 2004; García-Robledo & Horvitz 2012). In such cases there is no generalism-performance trade-off and specialization is a sub-optimal state for a consumer. The IHS was initially proposed as an explanation for this diversity of results.

The main question behind this dilemma is whether the same traits that allow a consumer species to efficiently exploit a given resource species do also allow it to exploit other resource species. This tends to be true if the resources are similar to one another, but false if not (Krasnov *et al.* 2004). Starting from this perspective, the IHS predicts that the relationship between consumer’s performance and specialization depends on resource heterogeneity. However, diverse communities can comprise clusters of similar resource species, each cluster being highly different from the other. For instance, the host community studied by Pinheiro *et al.* (2016) contains several birds species of the same genus, but also birds of different orders. In such cases of a wide range of resource dissimilarities, the IHS predicts a multi-scale relationship between performance and specialization. Considering only a group of similar resources, a “jack-of-all-trades” consumer tends to be master of all, though, between different clusters of resources the trade-off is strong (Pinheiro *et al.* 2016).

In previous studies, we proposed that the same mechanism governing the specialization *vs*. performance relationship may drive the architecture of consumer-resource networks (Pinheiro *et al.* 2016; Felix *et al.* 2017). From this perspective, nestedness is the result of the correlated performances of each consumer on similar resources, although modularity emerges because of strong trade-offs in performances on dissimilar resources. Therefore, the IHS predicts that subnetworks that represent phylogenetic or taxonomic subsets of complete systems, and thus do not comprise trade-offs, should be nested. However, in more diverse networks a multi-scale topology should emerge: a modular structure with internally nested modules.

This multi-scale architecture was named compound topology, a conceptual archetype proposed by Lewinsohn *et al.* (2006) and predicted by Flores *et al.* (2011). A compound topology is also a suitable explanation for networks that are nested and modular at the same time, because in those networks those conflicting topologies would predominate at different scales, instead of being mixed in the structure (as suggested by Fortuna *et al.* 2010). Evidence of a compound topology was found in pollination (Bezerra *et al.* 2009), bacteria-phage (Flores *et al.* 2013), and mammal-flea (Felix *et al.* 2017) empirical networks, as well as in synthetic networks (Beckett & Williams 2013; Leung & Weitz 2016). Moreover, a pattern of in-block nestedness was found in a large set of mutualistic and antagonistic networks, which, as far as we can tell, is the same structure as a compound topology (Solé-Ribalta *et al.* 2018).

Here, we propose a new mechanistic model for interaction networks based on the IHS. Our new model is presented in terms of consumers and resources, so it can help predict the topology of networks formed by different kinds of interaction, from antagonism to mutualism. The first-principles of our model are: (i) each resource species has a set of traits that affect its exploitability by each consumer species, and resource species can be more or less similar to one another in those traits; (ii) a consumer’s mutation that enhances its exploitation of a given resource tends to improve the exploitation of similar resources, but worsen its exploitation of dissimilar resources; and (iii) the capacity of a consumer to exploit each resource on a given moment is a result of its previous adaptations and maladaptations.

Following these simple principles, and adjusting a set of five parameters, we tested whether the IHS model can: (1) reproduce the varied relationships between performance and specialization of consumers observed in natural systems; (2) reproduce the main topologies observed in interaction networks, (3) explain the general conditions that affect the emergence of those patterns, and (4) generate predictions that are consistent with ecological and evolutionary theories and coherent with real-world observations. Moreover, our model is aimed to be a proof-of-concept (*sensu* Servedio *et al.* 2014) of the IHS, testing whether its predictions are logically derived from its assumptions and mechanism.

## The IHS model

### Core structure

Our model simulates the evolution of consumer species exploiting resource species. It is species-based and does not account for intraspecific variations. For increased text fluency, hereafter, we call consumer species “consumers” and resource species “resources”. Similarly, consumer species richness is referred to as “consumer richness”, and resource species richness as “resource richness”.

The core of our model consists of two evolving matrices: the innate performance matrix, and the realized performance matrix. In addition, there are two static inputs: a matrix with the pairwise distances between resources, and a vector of resource carrying capacities (Fig. 1).

**Figure 1.**
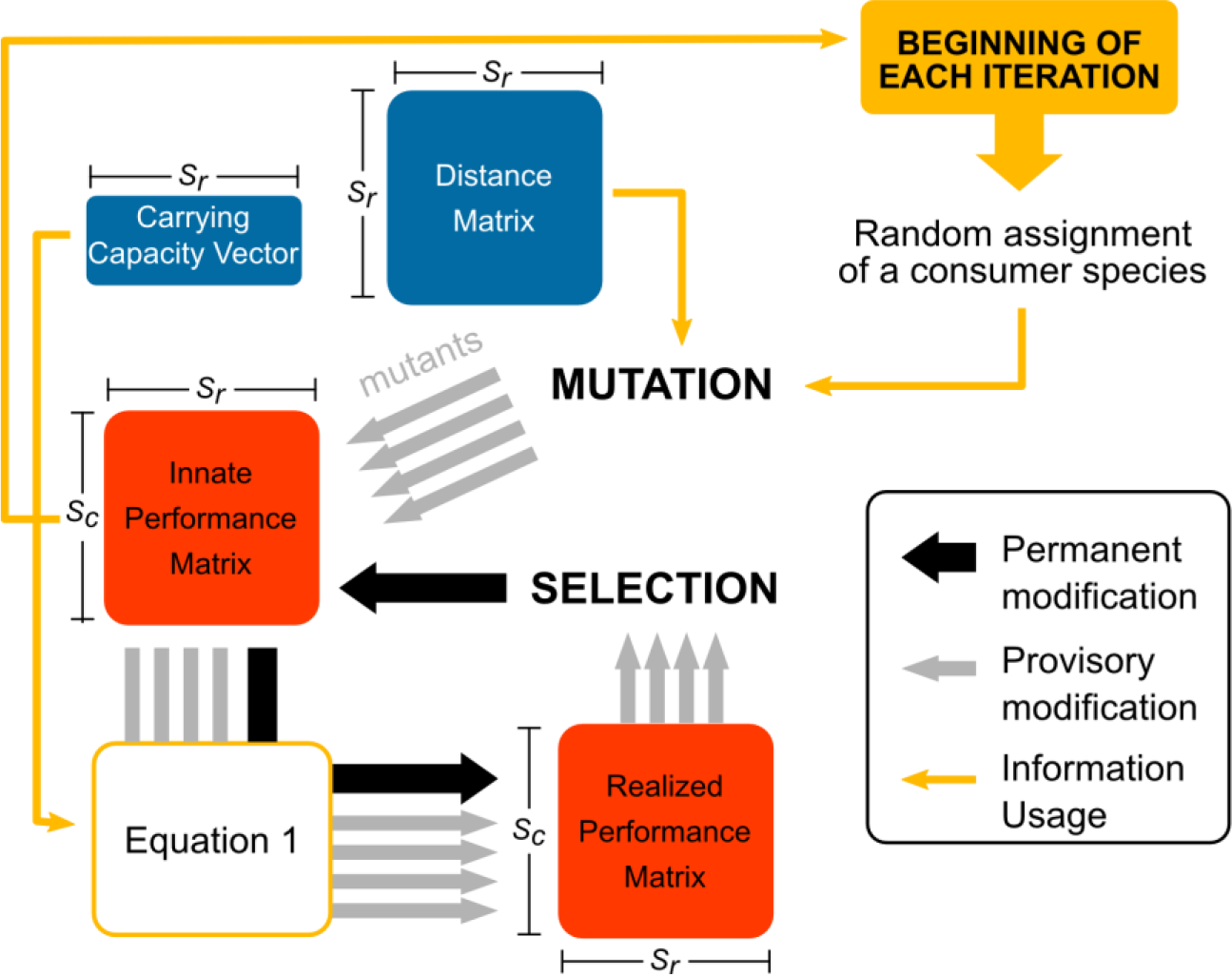
The IHS model. The iteration starts with the assignment of a random consumer that will evolve. This consumer suffers alternative mutations, each generating a mutant with its own innate performances on resources. Each mutation is focused on a given resource (focal resource) but affects the consumer’s innate performance on all resources. The consequence of each mutation for the consumer’s innate performance on a given resource depends on the distance between this resource and the focal resource, which is given by the resource species distance matrix. Then, using equation 1 (see the section “The IHS model”) the realized performance of each mutant is calculated. The mutant with the highest total realized performance is selected and replaces the original consumer in the innate performance matrix to be used in the next iteration of the model (unless all mutations result in decreased total realized performance, in which case the original consumer is maintained). For a detailed example of one iteration of the IHS model see Supplementary Figure S1. *S_c_*: consumer richness; *S_r_*: resource richness. Elements in blue are static inputs: do not change during the simulation. Elements in red are evolving matrices.

Innate performance represents the match between a consumer and a resource. It summarizes how all characteristics of the consumer (e.g., morphology, physiology, and behavior) affect its ability to exploit a given resource. When a consumer has a negative innate performance on a resource, it is incapable of exploiting it. However, when its innate performance is positive, the consumer exploits the resource (has a realized performance on it).

The distance between two resources in our model is a measure of how different they are from consumer’s perspective. Resources are close to one another when they require of the consumers the same adaptations for an efficient exploitation. For instance, two plant species, whose fruits have similar shape, size, and consistency, require from frugivorous birds the same type of beak. Resources are distant from one another when they require of the consumers opposite adaptations for an efficient exploitation. For instance, two plant species whose fruits are more easily consumed by, respectively, small-beaked and large-beaked birds. Because of phylogenetic conservatism, we expect the distances between resources to mirror the taxonomic and phylogenetic distances between them, however, convergence may confuse this pattern.

The carrying capacity of each resource limits the overall realized performance of its consumers. It can be understood as the availability of each resource for consumer exploitation. In natural systems, we expect abundance, size, and vulnerability (in antagonisms) or accessibility (in mutualisms) of each resource to be major factors defining this value.

Each realized performance represents the strength of an interaction effectively made in a consumer-resource system, therefore it cannot have a negative value. It integrates the match between consumers and resources (i.e., innate performance) with the limitations imposed by each resource carrying capacity, as presented in equation 1:

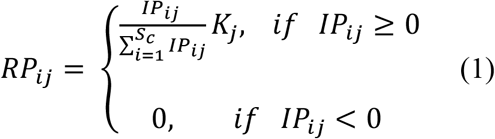

in which *IP*_*ij*_ is the innate performance of consumer *i* on resource *j*, *RP*_*ij*_ is the realized performance of consumer *i* on resource *j*, *S*_*c*_ is consumer richness, and *K*_*j*_ is the carrying capacity of resource *j*. In other words, consumers that have negative innate performances on a given resource, have zero realized performance on it. And for consumers that have positive innate performances on a given resource, the realized performances are the resource’s carrying capacity divided between these consumers proportionally to their innate performances.

### Mutation phase

At the beginning of each iteration, a consumer is randomly assigned to evolve. This consumer, then, is submitted to alternative mutations, one focused on each resource (focal resource), therefore generating *S*_*r*_ (resource richness) mutants of the consumer.

Mutations change the innate performance of the assigned consumer on all resources. The values of those changes are randomly drawn from normal distributions, in which standards deviations are equal to 0.3 and means are defined by the distance between each resource and the focal resource of the mutation, as presented in equation 2:

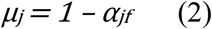

in which *μ*_*j*_ is the mean of the normal distribution from which we draw the value of changes in the innate performance of the assigned consumer on resource *j*, and *α*_*jf*_ is the distance between resource *j* and the focal resource *f*. Since the distance of the focal resource from itself is 0, the focal mutation will be a value randomly drawn from a normal distribution of mean = 1. Notice that, as a consequence of equation 3, each mutation probabilistically tends to increase the innate performance of the mutating consumer on resources with distances from the focal resource above 1 (*μ*_*j*_ > 0) and tends to decrease performances beyond this threshold (*μ*_*j*_ < 0).

### Selection phase

In the selection phase, following equation 1, the total realized performance of each mutant consumer is compared with the total realized performance of the original consumer (before mutations). If at least one mutant present increased total realized performance, the mutant with the largest total realized performance is selected, replacing the original consumer in the innate performance matrix for the next iteration (i.e., evolutionary changes occurred). However, if all mutations result in decreased total realized performance, the original consumer is selected, and the simulation goes to the next iteration without evolutionary changes.

### End of the simulation

The simulation ends after a pre-defined number of iterations. Then, by applying equation 1 on the final innate performance matrix, the final realized performance matrix is generated. This matrix corresponds to the simulated consumer-resource network (hereafter referred to as “simulated network”). Its contains the information concerning the consumer and resource species in the network (nodes), the interactions that are made between those species: consumers exploiting resources (links), and the consumers’ performance on exploiting each resource (weights). Moreover, as consumers cannot interact with other consumers, nor resources can interact with other resources, the simulated network is bipartite (two-mode).

For a complete example of an iteration of the IHS model, see Fig. S1 in Supporting Information.

## Simulations

### Inputs and parameters of the simulations

#### Innate performance matrix

To start each simulation, we need to provide an initial innate performance matrix. We built matrices with different consumer richness and resource richness. To fill the matrix we used three different methods: rep0) all consumers score 0 (zero) in innate performance on all resources, then the first mutation of a consumer corresponds to its ingress in the simulated network; rnorm11) the innate performance of each consumer on each resource is randomly drawn from a normal distribution with mean = 1 and standard deviation = 1; and rep1) all consumers score 1 (one) in innate performance on all resources.

#### Carrying capacity vector

The carrying capacity of each resource was defined by randomly drawing a value from a normal distribution with mean = 200 and standard deviation = 50.

#### Matrix of resource distances

The IHS predicts that network topology emerges as a function of the distance between resources and the degree of clustering of those distances. To test this prediction, we generated distance matrices defining values for the maximum distance between two resources and the number of clusters it contains (for details see Supplement S1).

#### Number of iterations

The number of iterations for each simulation was defined as each consumer has averaged 50 rounds of evolution. Therefore, the number of iterations equals consumer richness times 50.

#### List of parameters

In our simulations we adjusted five parameters: the consumer richness, the resource richness, the method used to generate the initial innate performance matrix (innate method), the maximum distance between two resources (maximum distance), and the number of resource clusters (number of clusters).

### Running simulations

Simulations were coded in R (R Core Team 2018). For commented codes see Supplement S1. The parameter values used in our simulations were: consumer richness: 5, 10, 50, 100, and 200; resource richness: 50, 100, and 200; innate method: rep0, rnorm11, and rep1; maximum distance: 1, 1.5, 2, 2.5, 3, 3.5, and 4; and number of clusters: 1, 2, and 4. We ran one simulation for each combination of those values, totalizing 945 setups.

## Statistical analysis

### Proportion of iterations in which occurred evolutionary changes

We used generalized linear models (GLM) to test which parameters affected the proportion of iterations in which occurred evolutionary changes in each simulation. In the complete model, we included as explanatory variables: (1) maximum distance, (2) innate method, (3) number of clusters (as a categorical variable), (4) resource richness, (5) consumer richness, and all interactions between variables (1), (2), and (3). After building the complete model, we used a backward stepwise approach with analysis of variance to reduce it to a minimum model. We used the explained deviance of each explanatory variable in the minimum model as a measure of effect size. This same setup was followed in all GLMs built in our study. For details about all statistical analyses performed in this study see Appendix S1.

For the subsequent analyses, we removed the simulations in which evolutionary changes occurred in less than 80% of iterations. There remained 672 simulations (72%).

### Relationship between performance and resource specialization of consumers

For each consumer in the simulated networks we calculated three performance indices: (1) mean realized performance, its average performance on all resources it exploits, (2) maximum realized performance, its maximum performance on a single resource, and (3) total realized performance, the sum of its performances on all resources. We also calculated two resource specialization indices, the first binary and the second weighted: (1) basic resource specialization, the richness of resource species exploited by the consumer, and (2) structural resource specialization, the diversity of resources exploited by the consumer measured with Shannon index (Poisot *et al.* 2012).

Then, we calculated Spearman correlations between the three performance indices and the two resource specialization indices for each simulated network. It was not possible to calculate the correlations using basic resource specialization for completely filled matrices, because in them, all consumers exploit the same resource richness.

To assess which factors influence the relationship between consumers’ performance and specialization in our simulations, we used generalized additive models (GAM) with the correlations as response variables and simulation parameters as explanatory variables. The maximum distance was included as a smooth term on each GAM. To find the minimum model we used the same approach used for the GLMs. In the present study, we used GAMs when the relationship between the response variable and maximum distance could not be properly modelled with a GLM.

### Network analysis

#### Network specialization

For each simulated network, we calculated a binary and a weighted network specialization metric: respectively, connectance and H_2_’ (Blüthgen *et al.* 2006). Connectance is defined as the proportion of potential links that are made in the network, therefore, the smaller its value, the more specialized the network. For H_2_’ the contrary is true: the higher it value, the more specialized the network. Specialization indices were computed using the package bipartite for R (Dormann *et al.* 2008). To test whether the simulation parameters influenced the specialization of the simulated networks we used GLMs.

#### Modularity

To measure the modularity and module composition of each simulated network we used the DIRTLPAwb+ algorithm (Beckett 2016), which maximizes the Barber modularity (Barber 2007) for weighted bipartite networks. Then we tested whether modularity values were affected by simulation parameters using GLMs.

#### Nestedness

To compute nestedness in weighted bipartite networks we used a new metric, which we named WNODA (weighted nestedness based on overlap and decreasing abundance). WNODA is a modification of WNODF (weighted nestedness based on overlap and decreasing fill) (Almeida-Neto & Ulrich 2011). WNODF is a nestedness metric designed for weighted networks, however, it maintains the condition of binary decreasing fill from the original NODF metric (Almeida-Neto *et al.* 2008). Therefore, WNODF can be strongly affected by weak links, which is not optimal for a weighted metric, and cannot deal with completely filled matrices (in those cases WNODF is 0). WNODA, in turn, does not demand binary nestedness to account for weighted nestedness, is less affected by weak links, and can be used for completely filled matrices. WNODA measures how frequently the weight of each link made by a node of lower total abundance is weaker than the weight of those same link made by a node with higher total abundance. Detailed information about WNODA and comparisons between metrics are presented in Appendix S2.

We calculated the WNODA of each simulated network and used GLMs to see how it was affected by the simulation parameters. To test the correlation between nestedness and modularity in our networks, we performed a Spearman correlation test.

Considering the possibility of a compound topology in our simulated networks, we used an approach based on the method proposed by Flores *et al.* (2013) and adapted by Felix *et al.* (2017), in which we separately compute the nestedness between species belonging to the same module and the nestedness between species belonging to different modules (Felix *et al.* 2017). This method can be performed with any nestedness metric based on pairwise comparison between nodes, including WNODA (see Appendix S2).

In a network with a compound topology we expect the WNODA between species of the same module (WNODA_SM_) to be much higher than the WNODA between species of different modules (WNODA_DM_). An R function to compute these components of nestedness using NODF, WNODF, and WNODA is provided in Supplement S2.

We used GLMs to test for effects of maximum distance and number of clusters on the WNODA_SM_ and WNODA_DM_ of the simulated networks.

#### Network topologies

In the present study, we considered three network topologies: modular, nested, and compound. To categorically define which topology was shown by each simulated network, we used the approach proposed by Felix *et al.* (2017) based on null model analysis.

First, we tested for nested and modular topologies using free null models. In the free models, each randomized matrix was generated using a modified version of the method proposed by Vázquez *et al.* (2007). Their method creates a null matrix conserving the original connectance and the total number of interactions, and probabilistically conserving the marginal sums. To this end, the algorithm first defines the binary structure of the null matrix, assigning interactions according to probabilities based on the marginal sums of the original matrix. However, to prevent reducing the size of the matrix, the algorithm requires that each species makes at least one interaction. After that, the remaining interactions are distributed among the filled cells, following again probabilities based on marginal sums. This method, however, is not fully adequate to our simulated matrices, as their interaction weights are not counts, but continuous. Therefore, the procedure results in null matrices with very different marginal sums from the original matrix, especially in matrices with many weak interactions. To deal with this, we modified the algorithm so that it does not fill the matrices by distributing unitary interactions (including and summing 1s) but by distributing a lower value. We defined this value as 0.1, as this was low enough to reasonable conserve the marginal sums.

For each simulated network, we generated a free null model with 500 randomized matrices and performed a Z-test to test whether the observed value of each metric was significantly different from the distribution of values of the null matrices. A network was considered modular when its value of Barber Modularity was significantly higher than the randomized values. Similarly, a network was considered nested, when it had a significant WNODA value. To avoid excessively low consumer richness in each module, we excluded the networks with 10 or fewer consumer species and kept 415 simulated networks for this and subsequent analysis.

A network was considered as having a compound topology, when it was significantly modular and presented a significant WNODA_SM_ (i.e., a modular network with modules internally nested). To test the significance of WNODA_SM_ in each simulated network we used restricted null models (Felix *et al.* 2017). A restricted null model is one that conserves the modular structure of the matrix when generating the randomized matrices. As, by definition, nodes in the same modules overlap more than nodes in different modules, not conserving the modular structure of the randomized matrix (i.e., using a free null model) would result in an inflated type I error ratio for WNODA_SM._

In the restricted null model, each interaction is first assigned an *a priori* probability and then the probabilities are adjusted to keep the modular structure. Here we used two different algorithms to assign the *a priori* probabilities of each interaction: Equiprobable and Degree-probable. In the Equiprobable method, *a priori* probabilities are equal for all cells and, therefore, only the modular structure defines the probability of each interaction. In the Degree-probable method, *a priori* probabilities are defined based on marginal sums (same method used for the free null model) and thenadjusted to maintain the modular structure of the matrix.

Null model analysis was performed in the Sagarana High-Performance Computing cluster from the High-Performance Processing Center, Institute of Biological Sciences, Federal University of Minas Gerais, Brazil.

We built GLMs to test how the simulation parameters affected the chance of a simulated network having a modular topology. Similarly, we tested for a nested topology. Then we tested, only for modular networks, how the simulation parameters affected their chances of having a compound topology.

### Multi-scale relationship between performance and specialization

To measure the resource specialization of consumers at different network scales, for each consumer in each modular network we calculated its standardized within-module degree (Z) and participation coefficient (P) (Guimerà & Nunes Amaral 2005). The first is a Z-score of the consumer’s degree within its module, and measures within-module specialization (small network scale). The second is a measure of how much the consumer’s links are distributed between different modules; therefore, it represents between-module specialization (large network scale). We also developed weighted versions of Z and P. The weighted Z is the Z-score of the diversity of links made by the consumer within its module, measured with Shannon index, and the weighted P measure the distribution of weights between modules.

As for the calculation of Z we need to compute standard deviations, it cannot be applied when all nodes of a module have the same degree. This resulted in some networks having too few usable values. For this analysis, we discarded networks with fewer than 5 nodes with meaningful values of both Z and P.

Then, for each network, we made linear regressions with consumer performances (mean, maximum and total) as response variables, and Z and P values (binary and weighted version) as explanatory variables. Finally, to test whether simulation parameters affected the relationship between performance and specialization of consumers at different network scales (i.e., coefficients of Z: β_Z_ and β_Weighted-Z_, and coefficients of P: β_Z_ and β_Weighted-Z_ in the linear regressions), we used GAMs.

## Results

The proportion of iterations in which occurred evolutionary changes decreased with maximum distance and number of clusters, and was lower in matrices built with the innate methods “rep1” and “rnorm11”. The other simulation parameters had low explanatory power (see Appendix S1.1). Out of the 945 simulations performed, 267 (28%) had less than 80% of the iterations with evolutionary changes and were removed from the subsequent analyses. The remaining simulations resulted in a highly diverse set of networks for every metric calculated in this study. Fig. 2 presents examples of this large variability.

**Figure 2.**
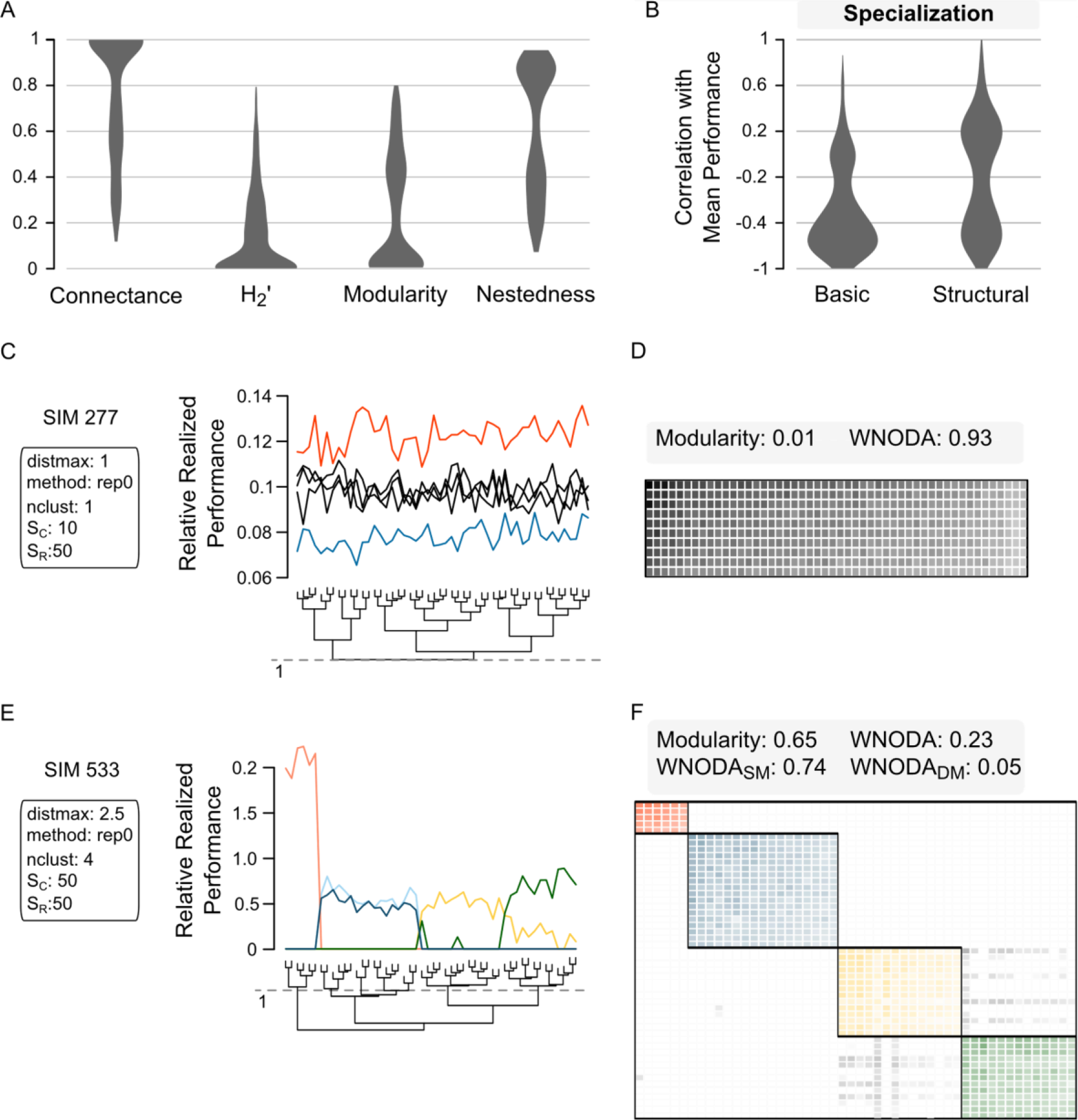
Diversity of patterns in the simulated networks. Our simulations resulted in a highly diverse set of consumer-resource networks considering all metrics analyzed (A). The relationship between specialization and performance of consumers varied largely (B). Here we illustrate two opposite patterns of specialization using as an example the simulated networks 277 (C-D) and 533 (E-F). In C and E, each line represents a consumer species. Five consumer species were sorted from each simulated system and their relative realized performances were plotted. The trees were obtained by hierarchical clustering of the distances between resources in the simulations, using the complete linkage method. Simulation 277 does not include performance trade-offs (maximum distance = 1) and does not have clusters in resource distance structure. The consumer which has the highest performance in one resource, also has the highest performances in all other resources (red line): the jack-of-all-trades is master of all. This simulation generated a network with very high nestedness and very low modularity (D). Rows and columns in D were organized by decreasing marginal sums and the grey tones represent the weight of each interaction. Nestedness is evidenced by the general trend of decreasing weights top-down and left-right in the matrix (D). Simulation 533 includes moderate trade-offs and clusters of similar resources. In this case, each consumer specializes in a group of similar resources (E). The network (F) has high modularity and low nestedness. Nevertheless, nestedness between species of the same module is high.

The correlation between mean performance and resource specialization depended on the distance between resources and the number of resource clusters, varying from positive to negative, and following the same general trend regardless of the resource specialization index used (Fig. 3A-C). The same trend held for the correlations with maximum performance (Appendix S1.2). The correlations involving total performance varied non-linearly with maximum distance. Our model predicts that specialists will present higher total performance than generalists when resources are intermediately distant one from another. Otherwise, generalists outperform specialists (Fig. 3D-E). See Appendix S1.2.

**Figure 3.**
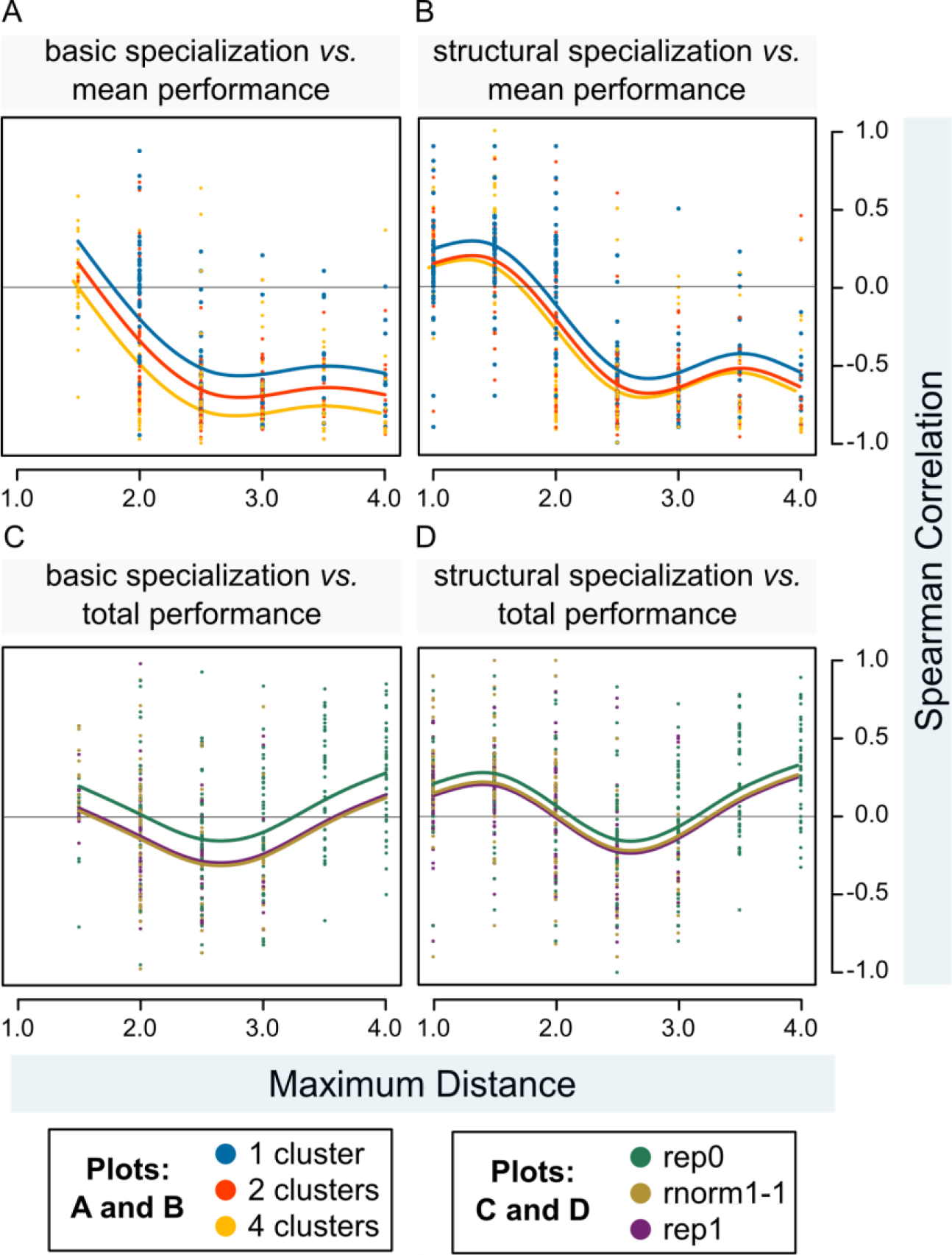
Correlations between performance and specialization of consumers. Correlations in the simulated network are presented as a function of maximum distance (horizontal axis), and number of clusters (colors in plots A and B) or innate method (colors in plots C and D). For each network we calculated Spearman correlations between indices of consumers’ realized performance (mean realized performance, maximum realized performance, and total realized performance) and indices of consumers’ specialization (basic specialization and structural specialization). Results for maximum performance were very similar to results for mean performance and are presented in Appendix S1. Notice that the values of specialization indices are negatively related to specialization, i.e., the higher the diversity of resources exploited by a consumer, the less specialized the consumer. The parameters represented in each plot are the ones with more explanatory power in the generalized additive models (see Appendix S1.2). In all plots, when consumer or resource richness were significant explanatory variables, we used its average values to draw the curves.

We found a consistent pattern of increasing network specialization with increasing maximum distance and number of clusters in simulations, in both the GLMs with connectance and H_2_’ (Fig. 4). Parameters related to the size of the network (consumer richness and resource richness) had just minor effects on connectance, but consumer richness had a moderate effect on H_2_’. Although the innate method defines the specialization of the initial matrix, it had little effect on connectance (Appendix S1.3) and H_2_’ (Appendix S1.4) in the simulated networks.

**Figure 4.**
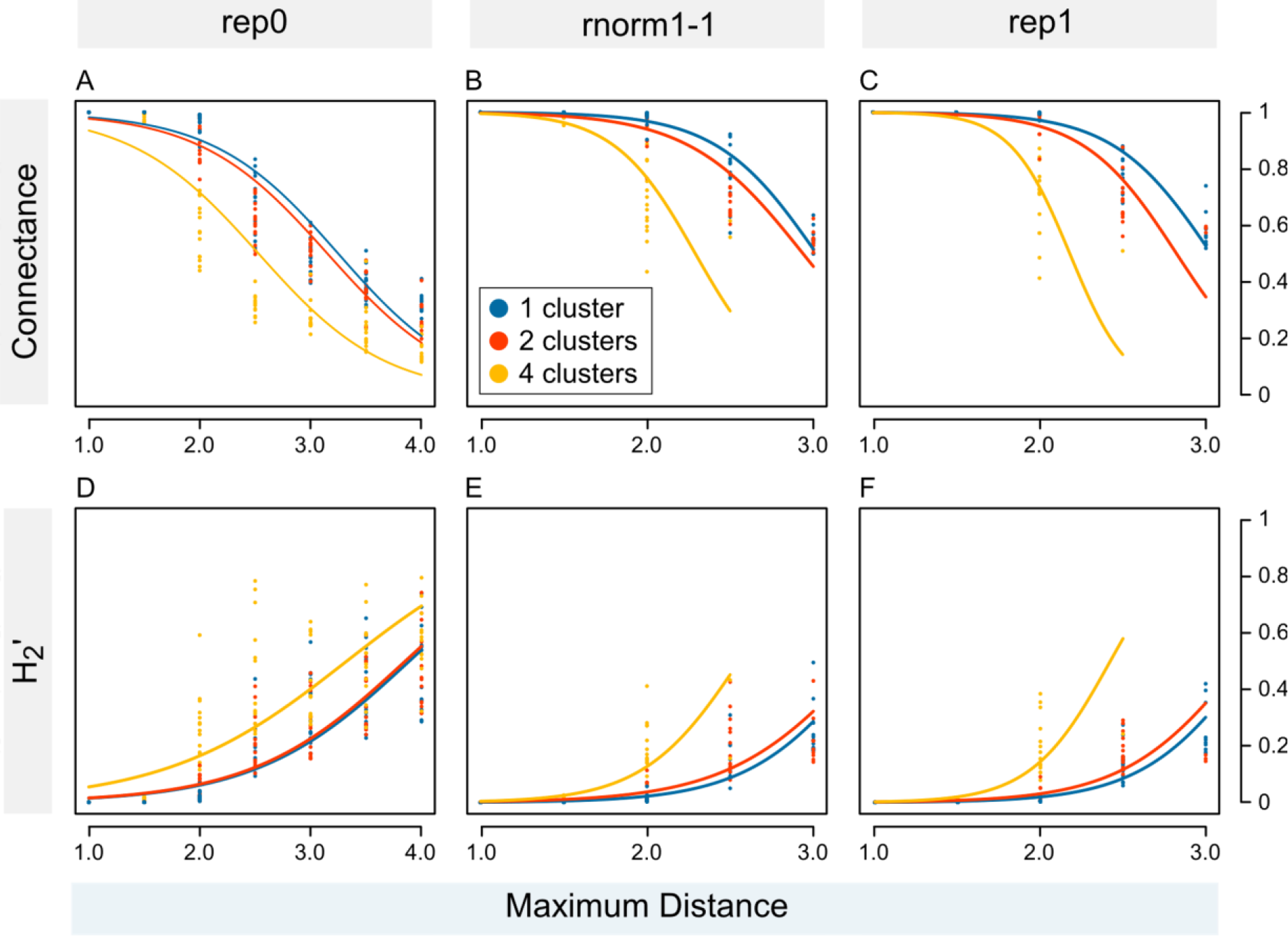
Network specialization metrics. Connectance and H_2_’ are presented as a function of maximum distance (horizontal axis), number of clusters (colors), and innate method (columns of plots in the panel). The parameters represented are the ones with more explanatory power in the generalized additive models (see Appendix S1.3-4). Average values of consumer and resource species richness were used to draw the curves. Notice that plots are presented with different scales in the horizontal axis.

Modularity increased with maximum distance and number of clusters (Fig. 5A), while nestedness decreased with those parameters (Fig. 5B). The other parameters had little or no effect on nestedness (Appendix S1.5) and modularity (Appendix S1.6) in the simulated networks. Both WNODA_SM_ and WNODA_DM_ decreased with maximum distance and number of clusters (Fig. 5C, Appendix S1.7). However, the former has a smaller slope than the later, and, therefore, the expected ratio between WNODA_SM_ and WNODA_DM_ increased with maximum distance and number of clusters (Fig. 5D). There is a strong negative correlation between modularity and nestedness on the simulated networks (Spearman rho: −0.94, p<0.001) (Fig. 5E, Appendix S1.8).

**Figure 5.**
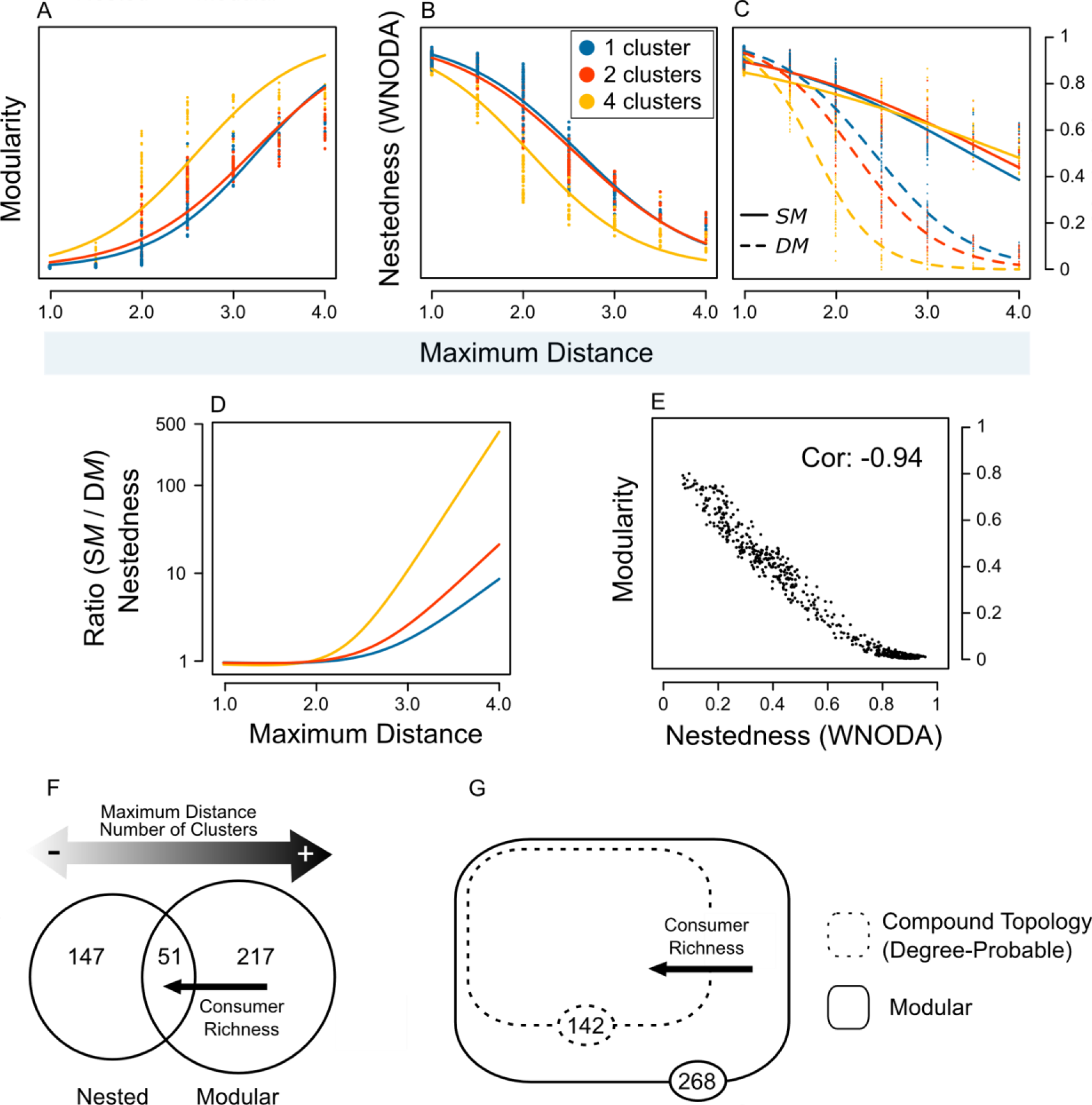
Simulation parameters affecting network topology. (A) and (B) show the effect of maximum distance and number of clusters on modularity and nestedness of the simulated networks, respectively. Values of WNODA were divided by 100, resulting in values between 0 and 1. In (C) nestedness is decomposed in its two components: nestedness between nodes of the same module (WNODA_SM_, solid lines) and nestedness between nodes of different modules (WNODA_DM_, dashed lines). Average values of consumer and resource richness were used to draw the curves in (A-C). Plot (D) shows the ratio between the expected WNODA_SM_ and WNODA_DM_ (curves on C) as a function of maximum distance. Notice that the vertical axis in D is log-transformed. In (E) a plot with nestedness *vs*. modularity shows the strong negative correlation between those metrics (Spearman correlation presented). (F) and (G) present Venn diagrams for network topologies and arrows for the main factors affecting the chance of a network showing each topology. The networks were classified as nested or modular based on null model analysis. Maximum distance and number of clusters have opposite effects on the chance of a network being nested or modular. Consumer richness had a strong effect on the probability of a network being nested, but a weak effect on its probability of being modular (F). Therefore, modular networks with high consumer richness have higher chance of also being nested. We tested whether each modular network presented a compound topology using restricted null models with two different methods to define *a priori* probabilities: equiprobable and degree-probable. All modular networks were detected as having compound topologies by the equiprobable restricted null model (not shown in the figure) and 142 were detected as having compound topologies by the degree-probable restricted null model (G). In this latter case, consumer species richness was the main factor influencing the probability of a modular network having internally nested modules (compound topology). All results presented here were obtained by fitting generalized linear models, except for (E), which was based on a Spearman correlation. Complete results are presented in Appendix S1.

From the 415 tested networks, 268 were significantly modular, 198 were significantly nested, and 51 were both modular and nested. The probability of a network having a modular topology increased with maximum distance and number of clusters, although the chance of a network being nested is affected by both parameters on the opposite direction. High consumer richness increased the chance of a simulated network being nested, but had a minor effect on the chance of it being modular. The other parameters had small effects on the models (Fig. 5F). Using the Equiprobable algorithm to define the *a priori* probabilities in the restricted null models, we detected that all modular networks showed in fact a compound topology. However, when the *a priori* probabilities were based on node degrees (Degree-probable), from the 268 modular networks, 142 were detected as having a compound topology. Using this last method, the main factor affecting the chance of a modular network presenting a compound topology was consumer richness. (Fig. 5G). For details see Appendix S1.9.

Most values of the β_P_ and β_Weighted-P_ in the regressions with mean performance were negative. However, this was not a ubiquitous pattern, as several positive values were also found. For Z the results were still more diverse, since most of the β_Z_ values were negative, although most of the β_Weighted-Z_ values were positive (Fig. 6A-B). In general, we found that the relationship between mean performance and Z decreased with maximum distance and number of clusters (Fig. 6D-F), although the relationship between mean performance and P was little or not affected by these parameters (Fig. G-I). The same general trends were found in the analysis using maximum performance instead of mean performance (Appendix S1.10). Similarly, most of the β_P_ and β_Weighted-P_ values in the regressions with total performance were also negative, and β_Weighted-Z_ values decreased with maximum distance, although this relationship was not observed for β_Z_ (Appendix S1.10).

**Figure 6.**
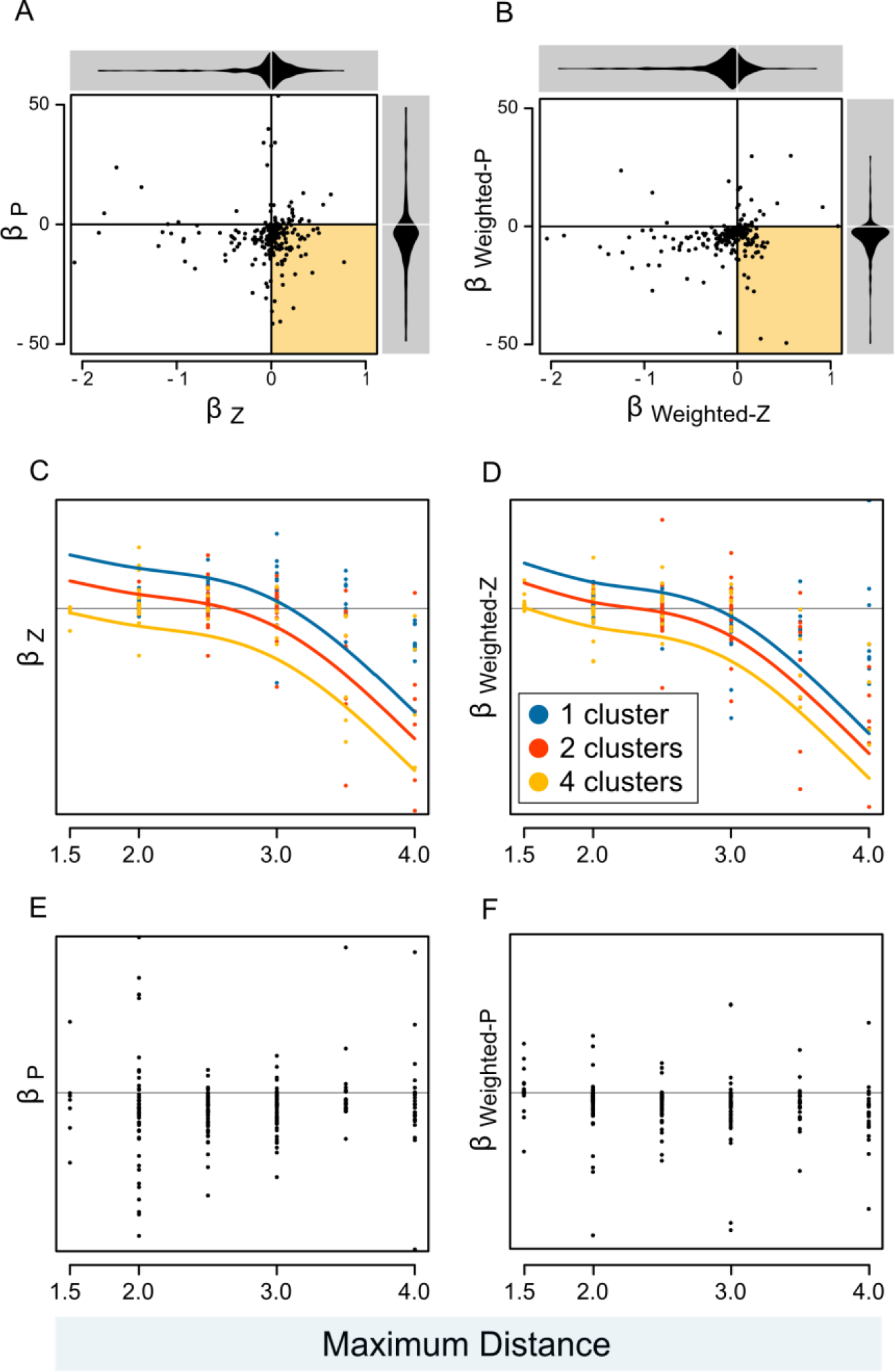
Simulation parameters affecting the multi-scale relationship between consumer’s mean performance and specialization. First, for each network we performed a linear regression between consumers’ mean performance as a function of Z (within-module degree) and P (participation coefficient). In (A) we plotted the coefficients (β) of these regressions. We also performed this same procedure using weighted versions of Z and P (B). The colored region of (A) and (B) represents the multi-scale relationship between performance and specialization predicted by the IHS: negative within-module (βZ>0) and positive between-modules (βP<0). Notice that the values of Z and P are negatively related to specialization. We built generalized additive models to test for a relationship between regression coefficients and simulation parameters (Appendix S1). In (C-F) we present the regression coefficients as a function of maximum distance (horizontal axis) and number of clusters (colors) when it has a statistically significant effect on the model. There were some coefficients with extreme values, whose inclusion would make it difficult to visualize the plots, and so, we show only the core region of each plot including most of the points and the predicted curves. Average values of consumer and resource richness were used to draw the curves.

## Discussion

The IHS model, following three first-principles, and through the adjustment of five biologically meaningful parameters, has successfully produced a highly diverse set of synthetic consumer-resource networks. In those simulations, specialization varied largely, and we found the main topologies observed in real-world interaction networks: nested, modular, and compound. We also found positive, neutral, and negative relationships between consumers’ performance and specialization, as well as multi-scale relationships. Despite this not being the first theoretical model to produce or predict one of those features separately (e.g., modularity: Guimerà *et al.* 2010; compound topology: Leung & Weitz 2016; positive relationship between performance and specialization: trade-off hypothesis, Poulin 1998; negative relationship between performance and specialization: resource breadth hypothesis, Hellgren *et al.* 2009), as far as we know, our model is the first to implement a single mechanism able to generate all patterns under different circumstances.

It is important to notice that no network-level structure was imposed on our model or emerged through network-level selection, but rather emerged from the rules on the evolution of links between consumers and resources. Moreover, by comparing simulated networks generated with different parameter setups we were able to identify general contexts that are related to the emergence of each pattern.

### Model parameters and simulated networks

Out of the five model parameters, maximum distance and number of clusters have disproportionately affected the simulated networks. Maximum distance is linked to the existence and intensity of trade-offs in consumer performances on different resources and number of clusters affects how discontinuously are those trade-offs distributed in the resource community. We found that discontinuities tend to reinforce the effect of increasing trade-offs on network architecture (i.e. maximum distance and number of clusters usually affect metrics of the simulated networks in the same direction).

The innate method defines the initial state of the network (the realized performance matrix before the simulation), however it had weak effect in most of the analysis of simulated networks (the realized performance matrix after the simulation), which shows that consumer evolution was strong enough to overcome initial patterns in most simulations. The only metric that was strongly driven by innate method was the proportion of iterations in which occurred evolutionary changes (for a discussion of this result, see Appendix S1.1). Overall, consumer and resource richness did not strongly influence the simulation outputs either, being important just in some analyses (e.g., compound topology), which we discuss further.

When using the IHS model, it is imperative to keep in mind that the simulated networks are ideal networks and several weak links on the matrices may not be detected in empirical studies or even may not happen in nature. First, it is well recognized that weak interactions are unlikely to being sampled in ecological studies (Jordano 2016). Also, some links may be so weak that it does not happen even once in ecological time or the resource exploitation is avoided by the consumer because it does not compensate the energy costs. For last, in most of interaction networks, weights are measured as counts (e.g. abundance of parasites in hosts, floral visits of pollinators), thus, imposing a lower limit on link weights (a link lower than 1 cannot occur). Therefore, despite several simulated networks have very high connectance, this is not likely to be found in empirical studies.

### Trade-offs and specialization

In general, higher values of maximum distance and number of clusters resulted in specialist consumers having higher performance than generalists on each resource, and in more specialized, more modular and less nested simulated networks. When trade-offs are strong, the jack-of-all-trades is master of none or, even, does not exist, and the network is sparse, with several forbidden links. However, when trade-offs are weak, the jack-of-all-trades is master of all, and the network is highly connected. When there is no trade-off at all (no distance between resources greater than 1), there is no forbidden links and connectance is always 1.

In natural systems we may expect that the intensity of trade-offs depends on the type and intimacy of the studied ecological interaction. As more intimate interactions require stronger match between interacting species than less intimate interactions (Hembry *et al.* 2018), the same difference between two resource species, tends to represent a stronger trade-off in intimate networks. For instance, slight physiological differences between two resources may strongly affect the probability of each resource being exploited by a given endoparasite, but be irrelevant to their probabilities of being preyed upon. In agreement with our predictions, ecological interactions known to be more intimate usually are more specialized than less intimate interactions (e.g., pollination *vs*. seed dispersal, Blüthgen *et al.* 2007; parasitism and parasitoidism *vs*. predatism, Van Veen *et al.* 2008; leaf mining *vs*. leaf chewing, Novotny *et al.* 2010), and form sparsely connected and modular networks, although low intimacy leads to highly connected and nested networks (Guimarães *et al.* 2007a; Pires & Guimaraes 2012; Hembry *et al.* 2018).

One of the most pervasive patterns in ecological networks is a negative relationship between size and connectance (Jordano 1987; Blüthgen *et al.* 2007). However, in our simulated networks, connectance was just minimally affected by consumer and resource richness (i.e., network size), but mostly driven by the intensity of trade-offs. Our results suggest that connectance in real-world ecological networks is not directly related to network size, but a consequence of larger networks usually comprising a more heterogeneous set of organisms and, therefore, containing stronger trade-offs. The same may hold for other network features that are affected by the intensity of trade-offs, such as modularity and nestedness. Using a similar rationale, Jordano (1987) argues that larger seed-dispersal networks are more compartmentalized and less connected because they include more diverse sets of feeding structures and fruit types. Moreover, Flores *et al.* (2011), in a set of nested networks, did not find a relationship between connectance and size, and Montoya *et al.* (2015) found that modularity in island networks was related with functional diversity but not with species richness, both corroborating that specialization decreases with heterogeneity and not with size itself.

### Compound topology

On the one hand, several simulated networks presented both significant nestedness and modularity. On the other hand, nestedness and modularity are driven in opposite directions by the same main parameters (maximum distance and number of clusters) and are strongly negatively correlated, as usually found in empirical ecological networks (Thebault & Fontaine 2010; Pires & Guimaraes 2012; Trøjelsgaard & Olesen 2013). This scenario does not support the perspective of the overall network having a mixed nested and modular structure (Fortuna *et al.* 2010), but is consonant with the perspective that each topology may predominate at different network scales (Felix *et al.* 2017).

Indeed, in modular-nested simulated networks, most of network nestedness came from the smaller scale: WNODA_SM_ was always much higher than WNODA_DM_. Our model predicts that networks without trade-offs should present a nested topology, and reinforces the prediction that highly diverse networks tend to present a compound topology (Lewinsohn *et al.* 2006; Flores *et al.* 2011; Felix *et al.* 2017). In these networks, consumers tend to specialize in a group of homogeneous resource species instead of a single species (Fig. 2D), which corroborates that network modules may be the real unity of specialization and coevolution (Olesen *et al.* 2007). Recently, as a result of conceptual and methodological improvements in ecological network science, compound topologies have been detected in several real-world networks that could be previously classified as purely modular or nested-modular (Flores *et al.* 2013; Felix *et al.* 2017; Solé-Ribalta *et al.* 2018).

We did not find a ubiquitous multi-scale relationship between consumer performance and specialization in modular networks, which suggests that this previously predicted pattern (Pinheiro *et al.* 2016) that has already been observed in nature (Felix *et al.* 2017), is not a necessary consequence of the IHS, but one of the possible structures that may emerge in diverse consumer-resource interaction systems. When the trade-offs are too strong, a positive relationship between performance and specialization emerges even within modules, which leads to extreme specialization. These are the situations in which we should expect to find pairwise specialization and coevolution. The relationship between performance and specialization in different modules presented a more random variation, that could not be explained based on the intensity of trade-offs in the simulations. If, on the one hand, a multi-scale relationship in modular networks was found in just a few cases, on the other hand, when entire simulated networks with increasing resource diversity are compared to one another, there is a clear inversion in the expected relationship between consumer performance and specialization.

Our results show that scale is key to understand the architecture and dynamics of ecological networks. And by scale we mean the hierarchical levels within a given network (e.g., network, modules, nodes), and the different taxonomic, phylogenetic, functional, and geographic scales that may be sampled when building a network from empirical data. Interaction networks containing only similar species show patterns that are not observed in more heterogeneous networks, as well as a module does not reflect the structure of the entire network. And, as previously commented by other authors, studies of ecological interactions are usually focused on modules of the network or in taxonomically defined assemblages subsets (Olesen *et al.* 2007; Jordano 2016). Thus, the literature is probably biased towards low-scale patterns (as suggested by Bezerra *et al.* 2009; Mello *et al.* 2011). This may explain, for instance, the paradigm of mutualism being always nested (Bascompte & Jordano 2007) and the dominance of positive relationships between performance and host range of parasites in the literature (Krasnov *et al.* 2004; Hellgren *et al.* 2009). Moreover, we may expect that several of the published nested interaction networks are in fact modules of more complete networks with compound topologies.

### Competing models that produce compound topologies

Beckett & Williams (2013) have predicted a compound topology for phage-bacteria networks, using a relaxed lock-and-key model. Despite their model including a larger number of parameters and having a more complex and less general mechanism than ours, most principles of the IHS model are at least partially met by it. In fact, only the first-principle (iii) of our model is not mirrored in some extent by their model, since performance is not defined only by consumer evolution, but also by resource evolution. We believe that our model is not contradictory to the relaxed lock-and-key model, but rather more comprehensive. Generality gets more and more important in those models, as observations of compound topologies in other systems are made (Felix *et al.* 2017; Solé-Ribalta *et al.* 2018).

Leung & Weitz (2016) proposed a bipartite network growth model that can also produce modular, nested, and compound networks. The mechanics of their model is very different from ours, mainly in two major aspects. First, in their model, a network grows by duplication of nodes, while in our model the number of species in the system is kept constant. Second, in their model once a link is stablished between two nodes it is not modified anymore, while in our model links depends on the match between consumers and resources, which is subjected to evolution. Moreover, contrary to the IHS model, their model produces only binary networks. These differences make it very difficult to compare assumptions and mechanisms of both models. However, it is remarkable that Leung & Weitz (2016) found that, when there are trade-offs, modularity emerges in networks, otherwise, hosts and parasites enter an arms race that results in nestedness. This is highly consonant with our main predictions using the IHS model.

### Limitations of the model

The main limitation of the IHS model is the assumption that innate performance is modified only by the evolution of consumer species. In nature, consumption is likely to be a selective force that also drives resource species evolution (Thompson 1994). This limitation is especially important in mutualisms, where it is not trivial to classify each partner as consumer or resource. In these cases, application of our model should take into account the available knowledge about the evolution of the species groups involved in the interaction. For instance, in pollination systems we may be eager to classify animals as consumers and plants as resources, because of the trophic relationship between them. However, there is strong evidence that plants evolve in response to pollinator-mediated selection, although the opposite is seldom true (Armbruster 2017). Therefore, it may be more appropriate to consider pollinators as resources exploited by plants in order to reproduce.

Another relevant limitation of our model is that the realized performance is determined only by resource species carrying capacity and innate performance, and does not consider consumer species abundance. This is a direct consequence of the IHS being initially proposed inspired by endoparasitic interactions. In obligatory interactions, from the consumers’ perspective, mainly when they are symbiotic, the abundance of the consumer species is itself a measure of interaction weight, as the consumer only survives by interacting. Then it is reasonable to consider consumer abundance and performance together in the model. However, in facultative interactions, in which consumer abundance is less dependent on the interaction, to not consider the separated effect of abundance and trait-matching in link establishment may represent a strong caveat.

Other limitations of the IHS model are: (1) the model does not include extinctions nor cladogenesis. It is important to warn that the present model does not aim to explain species coexistence in an ecological system but assumes it *a priori*. (2) The consumer-resource system is assumed to be closed: there is no emigration or immigration; and (3) links are affected just by the match between consumer and resource, overlooking factors exogenous to the species that may affect link establishment, e.g., context dependence (Chamberlain *et al.* 2014). Nevertheless, despite these somewhat simplistic assumptions, our model was able to recover all common topological patterns observed among interaction networks.

## Conclusion

In summary, we propose a new model for generating consumer-resource networks based on the integrative hypothesis of specialization (IHS). Despite its limitations, which are inherent to a model aiming at generality, our model may be a useful source of testable predictions.

One great challenge ahead is to parameterize our model based on real-world data, in order to generate more precise and quantitative predictions for particular kinds of networks. This is no simple task, though, as the distance between resource species is a non-dimensional variable, based on an abstract concept, which is affected by several factors. One possible solution would be to develop proxies for resource species distances based on phylogenetic, trait-based, or interaction-based distances.

However, even without these refinements, the proposed model reproduced several already observed patterns and most of its predictions are coherent to real-world observations and consonant with current evolutionary and ecological theories. Our results show that the IHS model is useful to generate synthetic, weighted, bipartite, consumer-resource networks and supports the IHS as a theoretical framework to study interaction specialization and network topology.

## Acknowledgements

We thank our institutions and many colleagues, who helped us in different ways during this project. We thank J. Miguel Ortega, Tetsu Sakamoto, and Verônica de Melo Costa for help with the use of Sagarana HPC cluster. The Graduate School in Ecology of the Federal University of Minas Gerais (ECMVS), Brazil, provided us with a scholarship from the Brazilian Council for Scientific and Technological Development (CNPq) granted to RBPP. Infrastructure for this study was provided by ECMVS, the Department of Ecology of the University of São Paulo (USP), and the Department of Biometry and Environmental System Analysis, University of Freiburg, Germany. The Graduate School in Ecology of the State University of Campinas (PPGE-UNICAMP), Brazil, provided GMFF with a scholarship from the Brazilian Coordination for the Improvement of Higher Education Personnel (CAPES). RBPP received a scholarship from the joint program between CAPES, CNPq, and Deutscher Akademischer Austauschdienst (DAAD) (88887.161398/2017-00) to make a one-year sandwich Ph.D at the University of Freiburg. MARM was funded by the Minas Gerais Research Foundation (FAPEMIG #PPM-00324-15), the Alexander von Humboldt Foundation (AvH: 3.4-8151/15037), and CNPq (#302700/2016-1).

## Authorship

All authors contributed to model development and study design. RBPP coded the model and performed the statistical analysis. All authors contributed to the interpretation of results. RBPP wrote the first draft and all authors reviewed the manuscript.

## Supporting Information

Figure S1 - An example of one iteration of the IHS model.

Supplement S1 - Code for the IHS model (ZIP file).

Supplement S2 - R function nest.smdm (ZIP file).

Appendix S1 - Supplementary analysis.

Appendix S2 - Weighted nestedness based on overlap and decreasing abundance (WNODA)

## References

Almeida-Neto, M., Guimarães, P., Guimarães, P.R., Loyola, R.D. & Ulrich, W. (2008). A consistent metric for nestedness analysis in ecological systems: reconciling concept and measurement. Oikos, 117, 1227–1239.

Almeida-Neto, M. & Ulrich, W. (2011). A straightforward computational approach for measuring nestedness using quantitative matrices. Environ. Model. Softw., 26, 173–178.

Armbruster, W.S. (2017). The specialization continuum in pollination systems: diversity of concepts and implications for ecology, evolution and conservation. Funct. Ecol., 31, 88–100.

Barber, M.J. (2007). Modularity and community detection in bipartite networks. Phys. Rev. E, 76, 066102.

Bascompte, J. & Jordano, P. (2007). Plant-animal mutualistic networks: the architecture of biodiversity. Annu. Rev. Ecol. Evol. Syst., 38, 567–593.

Bascompte, J., Jordano, P., Melian, C.J. & Olesen, J.M. (2003). The nested assembly of plant-animal mutualistic networks. Proc. Natl. Acad. Sci., 100, 9383–9387.

Beckett, S.J. (2016). Improved community detection in weighted bipartite networks. R. Soc. Open Sci., 3, 140536.

Beckett, S.J. & Williams, H.T.P. (2013). Coevolutionary diversification creates nested-modular structure in phage-bacteria interaction networks. Interface Focus, 3, 20130033–20130033.

Bellay, S., Lima, D.P., Takemoto, R.M. & Luque, J.L. (2011). A host-endoparasite network of Neotropical marine fish: are there organizational patterns? Parasitology, 138, 1945–1952.

Bezerra, E.L.S., Machado, I.C. & Mello, M.A.R. (2009). Pollination networks of oil-flowers: a tiny world within the smallest of all worlds. J. Anim. Ecol., 78, 1096–1101.

Blüthgen, N., Menzel, F. & Blüthgen, N. (2006). Measuring specialization in species interaction networks. BMC Ecol., 6, 9.

Blüthgen, N., Menzel, F., Hovestadt, T., Fiala, B. & Blüthgen, N. (2007). Specialization, constraints, and conflicting interests in mutualistic networks. Curr. Biol., 17, 341–346.

Chamberlain, S.A., Bronstein, J.L. & Rudgers, J.A. (2014). How context dependent are species interactions? Ecol. Lett., 17, 881–890.

Donatti, C.I., Guimarães, P.R., Galetti, M., Pizo, M.A., Marquitti, F.M.D. & Dirzo, R. (2011). Analysis of a hyper-diverse seed dispersal network: modularity and underlying mechanisms. Ecol. Lett., 14, 773–781.

Dormann, C.F., Fründ, J. & Schaefer, H.M. (2017). Identifying causes of patterns in ecological networks: opportunities and limitations. Annu. Rev. Ecol. Evol. Syst., 48, 559–584.

Dormann, C.F., Gruber, B. & Fründ, J. (2008). Introducing the bipartite Package: Analysing Ecological Networks. R News, 8, 8–11.

Felix, G.M., Pinheiro, R.B.P., Poulin, R., Krasnov, B.R. & Mello, M.A.R. (2017). The compound topology of a continent-wide interaction network explained by an integrative hypothesis of specialization. bioRxiv.

Flores, C.O., Meyer, J.R., Valverde, S., Farr, L. & Weitz, J.S. (2011). Statistical structure of host-phage interactions. Proc. Natl. Acad. Sci., 108, E288–E297.

Flores, C.O., Valverde, S. & Weitz, J.S. (2013). Multi-scale structure and geographic drivers of cross-infection within marine bacteria and phages. ISME J., 7, 520–532.

Fortuna, M.A., Stouffer, D.B., Olesen, J.M., Jordano, P., Mouillot, D., Krasnov, B.R., et al. (2010). Nestedness versus modularity in ecological networks: two sides of the same coin? J. Anim. Ecol., 79, 811–817.

Futuyma, D.J. & Moreno, G. (1988). The evolution of ecological specialization. Annu. Rev. Ecol. Syst., 19, 207–233.

García-Robledo, C. & Horvitz, C.C. (2012). Jack of all trades masters novel host plants: positive genetic correlations in specialist and generalist insect herbivores expanding their diets to novel hosts. J. Evol. Biol., 25, 38–53.

Guimarães, P.R., Rico-Gray, V., Oliveira, P.S., Izzo, T.J., dos Reis, S.F. & Thompson, J.N. (2007a). Interaction intimacy affects structure and coevolutionary dynamics in mutualistic networks. Curr. Biol., 17, 1797–1803.

Guimarães, P.R., Sazima, C., Reis, S.F. d. & Sazima, I. (2007b). The nested structure of marine cleaning symbiosis: is it like flowers and bees? Biol. Lett., 3, 51–54.

Guimerà, R. & Nunes Amaral, L.A. (2005). Functional cartography of complex metabolic networks. Nature, 433, 895–900.

Guimerà, R., Stouffer, D.B., Sales-Pardo, M., Leicht, E.A., Newman, M.E.J. & Amaral, L.A.N. (2010). Origin of compartmentalization in food webs. Ecology, 91, 2941–2951.

Hellgren, O., Pérez-Tris, J. & Bensch, S. (2009). A jack-of-all-trades and still a master of some: prevalence and host range in avian malaria and related blood parasites. Ecology, 90, 2840–2849.

Hembry, D.H., Raimundo, R.L.G., Newman, E.A., Atkinson, L., Guo, C., Guimarães P.R., et al. (2018). Does biological intimacy shape ecological network structure? A test using a brood pollination mutualism on continental and oceanic islands. J. Anim. Ecol., 0–2.

Ings, T.C., Montoya, J.M., Bascompte, J., Blüthgen, N., Brown, L., Dormann, C.F., et al. (2009). Ecological networks - Beyond food webs. J. Anim. Ecol., 78, 253–269.

Jordano, P. (1987). Patterns of mutualistic interactions in pollination and seed dispersal: connectance, dependence asymmetries, and coevolution. Am. Nat., 129, 657–677.

Jordano, P. (2016). Sampling networks of ecological interactions. Funct. Ecol., 30, 1883–1893.

Krasnov, B.R., Fortuna, M.A., Mouillot, D., Khokhlova, I.S., Shenbrot, G.I. & Poulin, R. (2012). Phylogenetic signal in module composition and species connectivity in compartmentalized host-parasite Networks. Am. Nat., 179, 501–511.

Krasnov, B.R., Poulin, R., Shenbrot, G.I., Mouillot, D. & Khokhlova, I.S. (2004). Ectoparasitic “jacks-of-all-trades”: relationship between abundance and host specificity in fleas (Siphonaptera) parasitic on small mammals. Am. Nat., 164, 506–516.

Krishna, A., Guimarães Jr, P.R., Jordano, P. & Bascompte, J. (2008). A neutral-niche theory of nestedness in mutualistic networks. Oikos, 117, 1609–1618.

Leung, C.Y. (Joey) & Weitz, J.S. (2016). Conflicting attachment and the growth of bipartite networks. Phys. Rev. E, 93, 032303.

Lewinsohn, T.M., Inácio Prado, P., Jordano, P., Bascompte, J. & M. Olesen, J. (2006). Structure in plant-animal interaction assemblages. Oikos, 113, 174–184.

Mello, M.A.R., Marquitti, F.M.D., Guimarães, P.R., Kalko, E.K.V., Jordano, P. & de Aguiar, M.A.M. (2011). The modularity of seed dispersal: differences in structure and robustness between bat–and bird–fruit networks. Oecologia, 167, 131–140.

Montoya, D., Yallop, M.L. & Memmott, J. (2015). Functional group diversity increases with modularity in complex food webs. Nat. Commun., 6, 7379.

Muchhala, N. (2007). Adaptive trade‐off in floral morphology mediates specialization for flowers pollinated by bats and hummingbirds. Am. Nat., 169, 494–504.

Novotny, V., Miller, S.E., Baje, L., Balagawi, S., Basset, Y., Cizek, L., et al. (2010). Guild-specific patterns of species richness and host specialization in plant-herbivore food webs from a tropical forest. J. Anim. Ecol., 79, 1193–1203.

Olesen, J.M., Bascompte, J., Dupont, Y.L. & Jordano, P. (2007). The modularity of pollination networks. Proc. Natl. Acad. Sci. U. S. A., 104, 19891–19896.

Pinheiro, R.B.P., Félix, G.M.F., Chaves, A. V., Lacorte, G.A., Santos, F.R., Braga, É.M., et al. (2016). Trade-offs and resource breadth processes as drivers of performance and specificity in a host–parasite system: a new integrative hypothesis. Int. J. Parasitol., 46, 115–121.

Pires, M.M. & Guimaraes, P.R. (2012). Interaction intimacy organizes networks of antagonistic interactions in different ways. J. R. Soc. Interface, 10, 20120649–20120649.

Poisot, T., Canard, E., Mouquet, N. & Hochberg, M.E. (2012). A comparative study of ecological specialization estimators. Methods Ecol. Evol., 3, 537–544.

Poulin, R. (1998). Large-scale patterns of host use by parasites of freshwater fishes. Ecol. Lett., 1, 118–128.

R Core Team (2017). R: A language and environment for statistical computing. R Foundation for Statistical Computing, Vienna, Austria. URL https://www.R-project.org/

Servedio, M.R., Brandvain, Y., Dhole, S., Fitzpatrick, C.L., Goldberg, E.E., Stern, C.A., et al. (2014). Not just a theory - The utility of mathematical models in evolutionary biology. PLoS Biol., 12, e1002017.

Solé-Ribalta, A., Tessone, C.J., Mariani, M.S. & Borge-Holthoefer, J. (2018). Revealing in-block nestedness: Detection and benchmarking. Phys. Rev. E, 97, 062302.

Stang, M., Klinkhamer, P.G.L. & van der Meijden, E. (2007). Asymmetric specialization and extinction risk in plant–flower visitor webs: a matter of morphology or abundance? Oecologia, 151, 442–453.

Thebault, E. & Fontaine, C. (2010). Stability of ecological communities and the architecture of mutualistic and trophic networks. Science (80-.)., 329, 853–856.

Thompson, J.N. (1994). The coevolutionary process. University of Chicago Press, Chicago, USA.

Trøjelsgaard, K. & Olesen, J.M. (2013). Macroecology of pollination networks. Glob. Ecol. Biogeogr., 22, 149–162.

Ulrich, W., Kryszewski, W., Sewerniak, P., Puchałka, R., Strona, G. & Gotelli, N.J. (2017). A comprehensive framework for the study of species co-occurrences, nestedness and turnover. Oikos, 1607–1616.

Vázquez, D.P., J. Melián, C., M. Williams, N., Blüthgen, N., R. Krasnov, B. & Poulin, R. (2007). Species abundance and asymmetric interaction strength in ecological networks. Oikos, 116, 1120–1127.

Van Veen, F.J.F., Müller, C.B., Pell, J.K. & Godfray, H.C.J. (2008). Food web structure of three guilds of natural enemies: predators, parasitoids and pathogens of aphids. J. Anim. Ecol., 77, 191–200.

Watts, S., Dormann, C.F., Martín González, A.M. & Ollerton, J. (2016). The influence of floral traits on specialization and modularity of plant–pollinator networks in a biodiversity hotspot in the Peruvian Andes. Ann. Bot., 118, 415–429.

